# Regulation of specific cell clusters in TCR-T cells responding to differential expression of tumor PD-L1

**DOI:** 10.1101/2020.07.28.224816

**Authors:** Renpeng Ding, Shang Liu, Huanyi Chen, Bin Kang, Radoje Drmanac, Ying Gu, Xuan Dong, Qianqian Gao

**Author notes:** These authors contributed equally (Renpeng Ding, Shang Liu). Correspondence: Qianqian Gao,; Xuan Dong,; Ying Gu, Radoje Drmanac,.

## Abstract

PD-L1 signaling is important in regulating T cell function and keeping balance of tumor microenvironment, but its role in modifying TCR-T cell cytotoxicity remains unknown. MART-1-specific TCR-T cells (TCR-T_MART-1_) were stimulated by MEL-526 tumor cells expressing different proportions of PD-L1 and used to perform cytotoxicity assays and single-cell RNA sequencing. Percentage changes of different specific cell clusters were analyzed. The percentage of cluster HLA-DR^+^CD38^+^CD8^+^ was upregulated after antigen stimulation and tumor PD-L1 modified TCR-T cell function through downregulating the percentages of clusters HLA-DR^+^CD28^+^CD8^+^ and HLA-DR^+^CD38^+^CD8^+^ which were higher in TCR-T_MART-1_ than in T_null_.

## Introduction

PD-L1 (programmed death-ligand 1) takes participation in regulating T cell-mediated immune responses for tumor evasion from the immune system, thus promotes cancer development and progression [1]. PD-L1 is also known as CD274 or B7-H1 and is one ligand for PD-1 (programmed death-1). PD-L1 is wildly expressed on tumor cells of various types of malignancies including melanoma, while PD-1 is highly expressed in tumor-infiltrating lymphocytes [2]. PD-L1 interacts with PD-1 resulting in T cell dysfunction and exhaustion, but the effect of tumor PD-L1 expression on TCR-T (T-cell receptor-engineered T cells) cell function has not been comprehensively studied. TCR-T cell therapy has great potential in mitigating tumor development especially for solid tumors. The number of clinical trials with TCR-T cell therapy is increasing each year and among them the most targeted cancer type is melanoma [3]. Therefore, it’s important to investigate how tumor PD-L1 expression affects TCR-T cell functionality.

In our study, single-cell mRNA sequencing (scRNA-seq) was performed to investigate MART-1-specific TCR-T cells responding to different proportions of PD-L1^+^ melanoma cells. Proportions of specific cell clusters such as HLA-DR^+^CD28^+^CD8^+^ and HLADR^+^CD38^+^CD8^+^ were modified with increasing ratio of tumor PD-L1.

## Results

### TCR-T_MART-I_ killed MEL-526 tumor cells efficiently at E:T ratio of 1:1

HLA-A*0201 /MART-1-specific TCR sequence was obtained from T cells after stimulation with MART-1 (aa27-35, LAGIGILTV) peptide (data unpublished) and designed as TCR_MART-1_ (Fig. 1). To avoid miss pairing with endogenous TCR [4], TCR_MART-1_ α and β chains were fused with constant region of murine TCR and synthesized before cloned into the lentiviral vector (Fig. 1). After transfection of TCR_MART-1_ lentivirus, CD8^+^ T cells expressed TCR_MART-1_ or not were designed as TCR-T_MART-1_ and T_null_, respectively. TCR-T_MART-1_ and T_null_ cells were stimulated with peptide-loaded MEL-526 melanoma cells at different E:T ratios (1:1, 1:2, and 1:4. Fig. 2) to assess the killing capacity. Compared to T_null_, TCR-T_MART-1_ killed tumor cells more efficiently especially at E:T ratio of 1:1 (Fig. 2).

**Figure 1.**
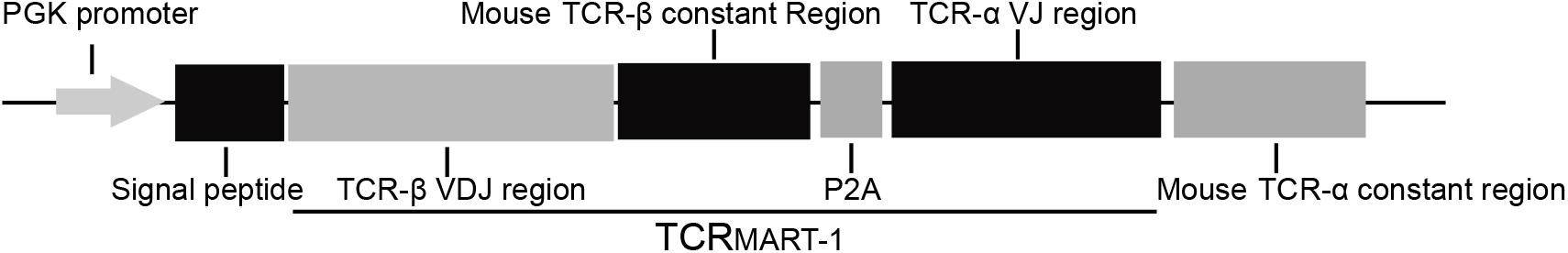
Schematic design of MART-1_26-35_ peptide-specific TCR sequence. The transcription of TCR_MART-1_ was driven by PGK promoter. Mouse constant regions were used to reduce the mispairing with endogenous TCR. TCRβ and TCRα were linked by P2A.

**Figure 2.**
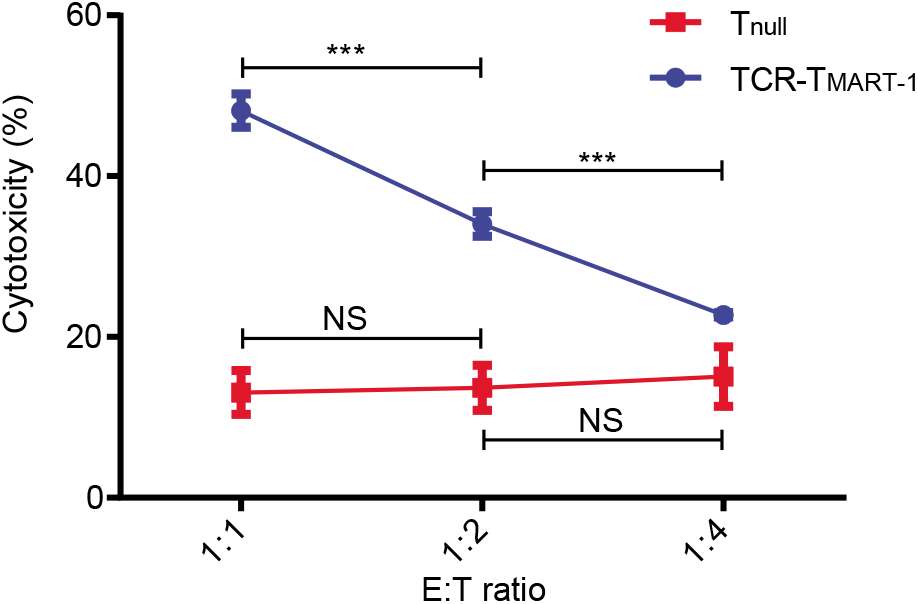
Cytotoxicity of T_null_ and TCR-T_MART-1_ against peptide-loaded MEL-526 cells. The in vitro killing assays were performed at different E:T ratios (1:1, 1:2, and 1:4). Data were generated from three individual replicates and shown as mean ± SD. 2-tailed unpaired t-tests were used to calculate p-values, *: p< 0.05; **: p < 0.01; ***: p < 0.001; NS: Not significant.

### Regulation of specific T cell clusters responding to different proportions of PD-L1^+^ tumor cells

To verify the effect of tumor PD-L1 expression on TCR-T cell function, MEL-526 cells expressing low, intermediate, and high levels of PD-L1 (data unpublished, designed as PD-L1_low_, PD-L1_int_, and PD-L 1_high_, respectively and PD-L1 expression ratio was about 3%, 50%, 100%) were incubated with TCR-T_MART-1_ (50% TCR_MART-1_^+^). The percentage of specific CD8^+^ T cell clusters including CX3CR^+^, HLA-DR^+^, HLA-DR^+^CD28^+^ and HLA-DR^+^CD38^+^, were analyzed in T cells (Fig. 3). The percentages of CX3CR^+^, HLA-DR^+^, and HLADR^+^CD28^+^ clusters were decreased, while the percentage of HLA-DR^+^CD38^+^ cluster was increased after antigen stimulation (Fig. 3). Furthermore, the percentages of HLA-DR^+^CD28^+^ and HLA-DR^+^CD38^+^ clusters were reduced by the increasing proportion of tumor PD-L1 (Fig. 3).

**Figure 3.**
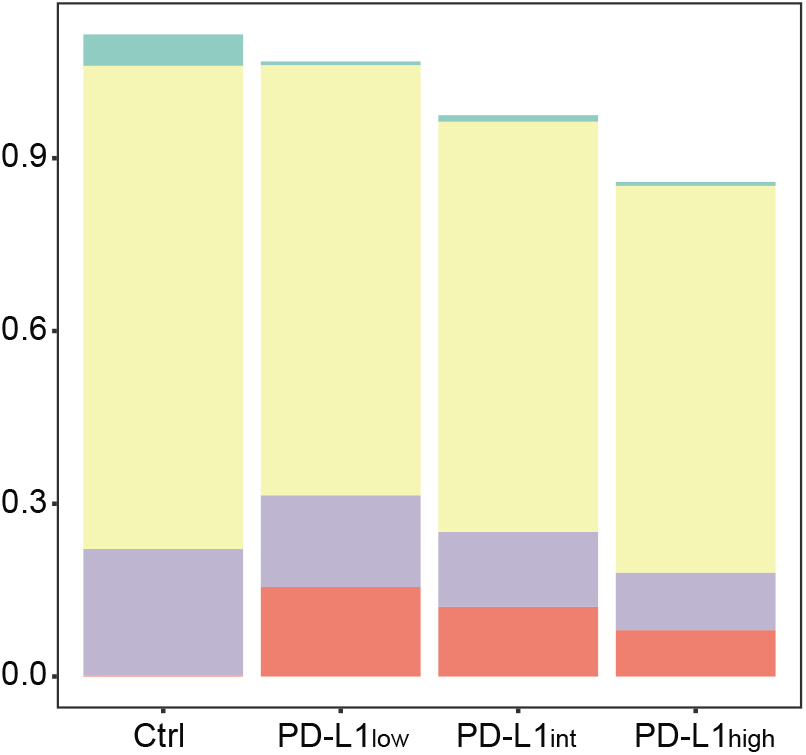
Compositions of specific cell clusters in T cells responding to tumor PD-L1. The proportions of clusters CX3CR1^+^, HLA-DR^+^, HLA-DR^+^CD28^+^, and HLA-DR^+^CD38^+^ in T cells stimulated with antigen or not.

When T cells were further divided into T_null_ and TCR-T_MART-1_, the percentages of HLA-DR^+^CD28^+^ and HLA-DR^+^CD38^+^ clusters were higher in TCR-T_MART-1_ than in T_null_ (Fig. 4). Consistently, the percentages of HLA-DR^+^CD28^+^ and HLA-DR^+^CD38^+^ clusters in both T_null_ and TCR-T_MART-1_ was downregulated with the increasing proportion of tumor PD-L1 (Fig. 4).

**Figure 4.**
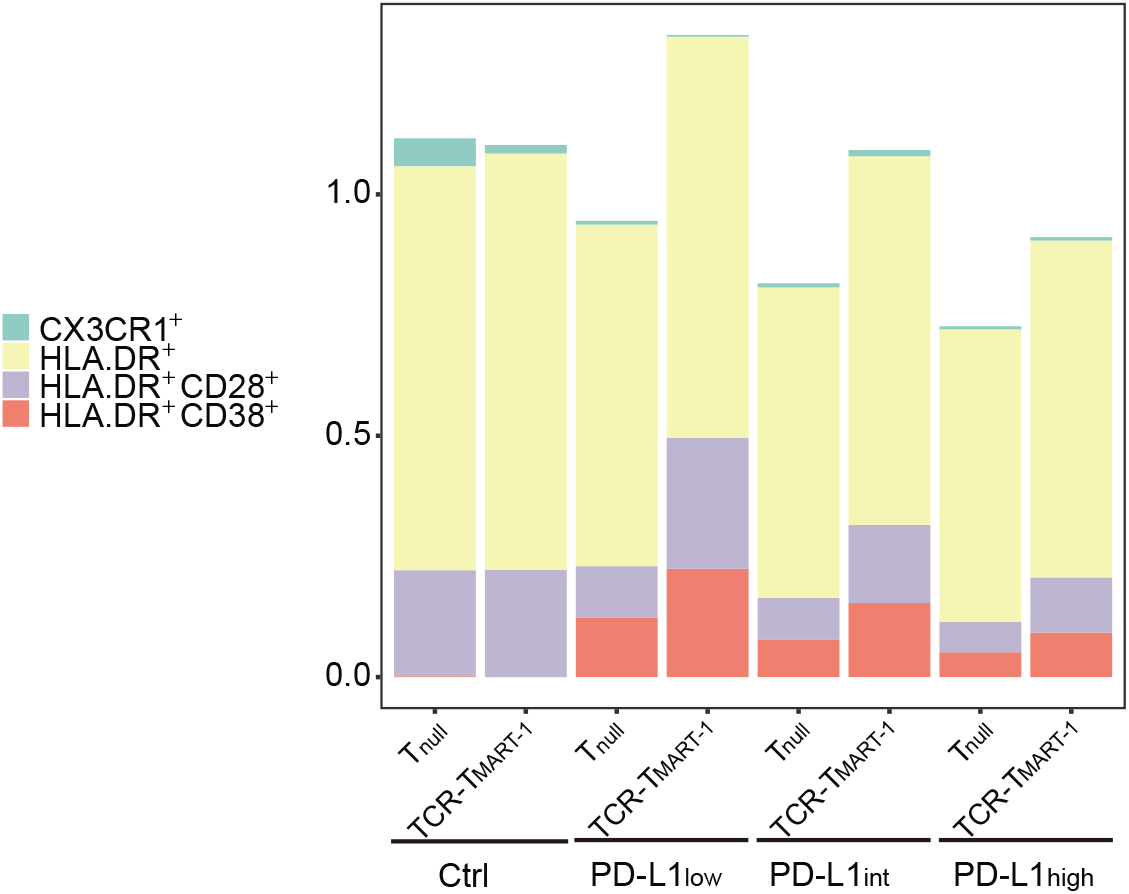
Compositions of specific cell clusters in T_null_ and TCR-T_MART-1_ responding to tumor PD-L1. The proportions of clusters CX3CR1^+^, HLA-DR^+^, HLA-DR^+^CD28^+^, and HLA-DR^+^CD38^+^ in T_null_ and TCR-T_MART-1_ stimulated with antigen or not.

### CX3CR^+^CD8^+^ cluster was characterized by *GZMA* expression

Differentially expressed genes (DEGs) were analyzed in these specific clusters. Except *CX3CR1,* the expression of cytotoxic genes *GZMA* and *NKG7,* and chemokine *CCL5* was upregulated in CX3CR^+^ cluster, while the expression of *IL2RA, XCL1,* and *GZMB* were downregulated compared to CX3CR^−^ cells (Fig. 5A). After gene oncology (GO) analysis, T cell activation and cell-cell adhesion related signaling were enriched in CX3CR^+^ cluster (Fig. 5B).

**Figure 5.**
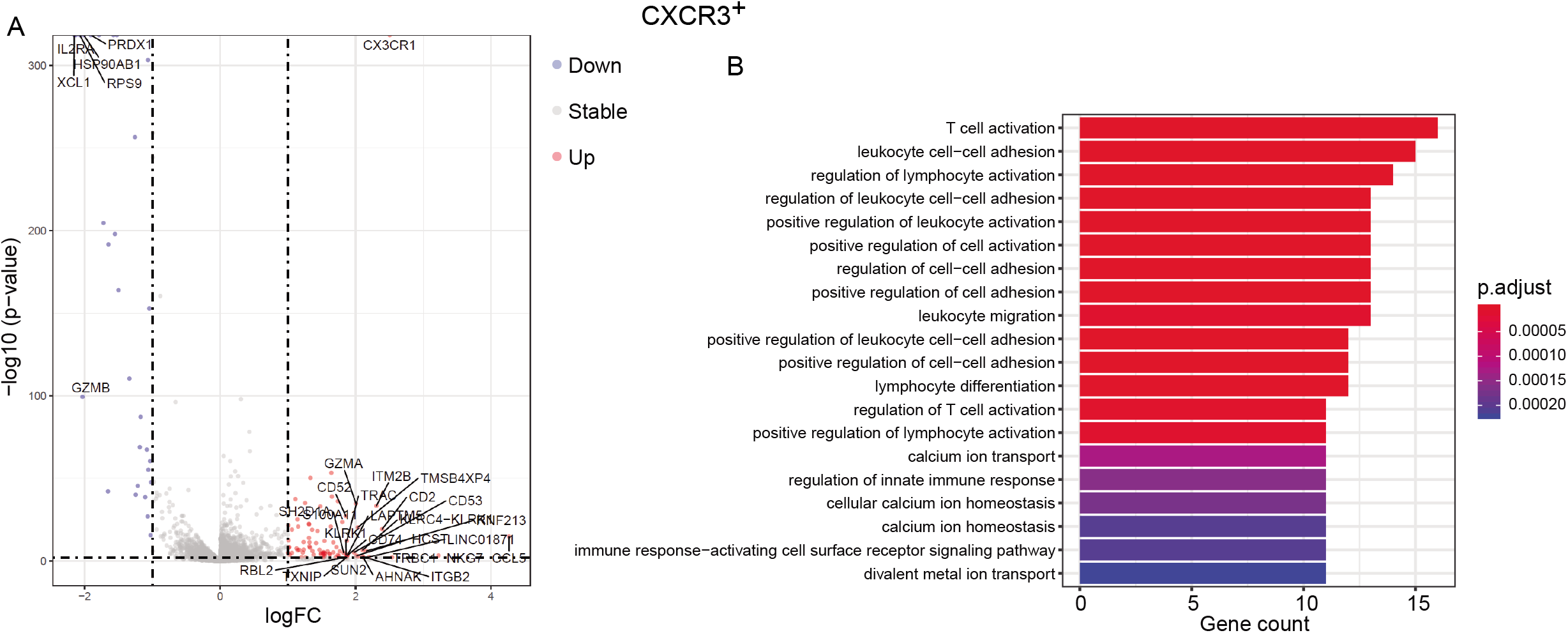
Characteristics of cluster CX3CR1^+^CD8^+^. A, Volcano plot showing differentially expressed genes between CX3CR1^+^ and CX3CR1^−^ T cells. Each red or blue dot denotes an individual upregulated or downregulated gene (|logFC| >= 1 and p.value < 0.01). B, Bar plot showing the top 20 pathways of CX3CR1^+^CD8^+^ T cells. The color represents p.value and the x-axis represents gene count.

### HLA-DR^+^CD8^+^ cluster was characterized by *IL32* and *GZMA* expression

In addition to *GZMA* and *CCL5* expression, which was upregulated in CX3CR^+^ cluster as well, the expression of cytokine *IL32* was increased in HLA-DR^+^CD8^+^ cluster (Fig. 6A). Endocytic vesicle membrane signaling were enriched in HLA-DR^+^CD8^+^ cluster (Fig. 6B).

**Figure 6.**
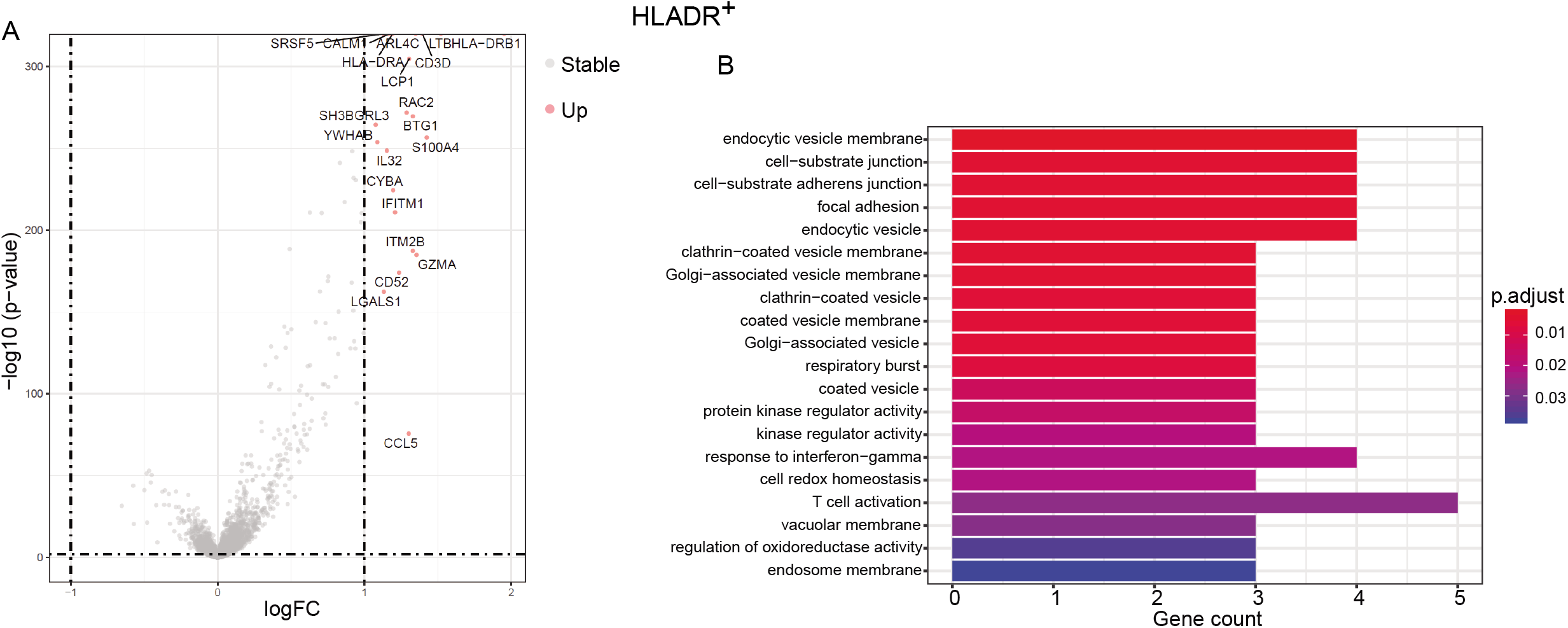
Characteristics of cluster HLA-DR^+^CD8^+^. A, Volcano plot showing differentially expressed genes between HLA-DR^+^ and HLA-DR^−^ T cells. Each red or blue dot denotes an individual upregulated or downregulated gene (|logFC| >= 1 and p.value < 0.01). B, Bar plot showing the top 20 pathways of HLA-DR^+^CD8^+^ T cells. The color represents p.value and the x-axis represents gene count.

### HLA-DR^+^CD28^+^CD8^+^ cluster was characterized by *CD52* expression

The expression of *CD52* was increased in addition to *CD28* in HLA-DR^+^CD28^+^CD8^+^ cluster (Fig. 7A). Though not so dramatic as that of *CD52,* the expression of *CCL5* and *JAK1,* which are essential for cytokine signaling, was upregulated as well. Leukocyte activation related pathways were enriched in this cluster (Fig. 7B), which was much similar with that of CX3CR^+^CD8^+^ cluster (Fig. 5B).

**Figure 7.**
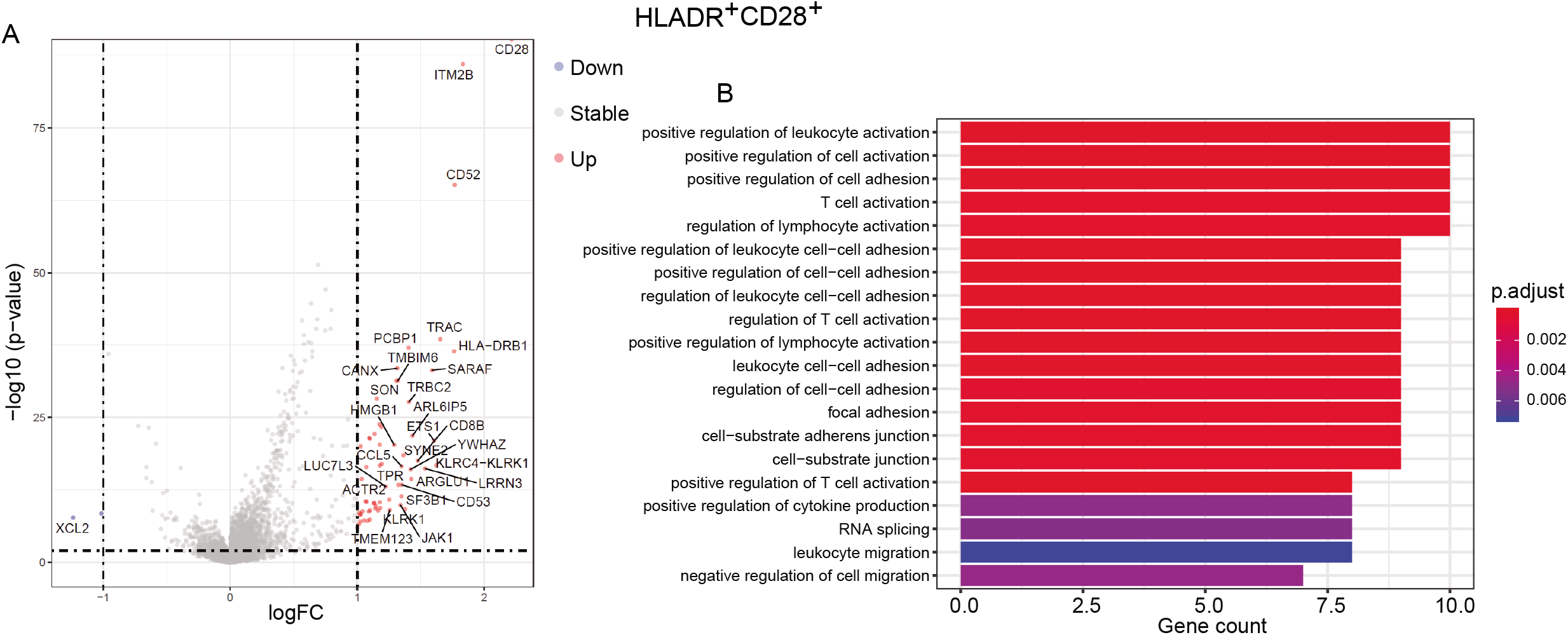
Characteristics of cluster HLA-DR^+^CD28^+^CD8^+^. A, Volcano plot showing differentially expressed genes between HLA-DR^+^CD28^+^ and non-HLA-DR^+^CD28^+^ T cells. Each red or blue dot denotes an individual upregulated or downregulated gene (|logFC| >= 1 and p.value < 0.01). B, Bar plot showing the top 20 pathways of HLA-DR^+^CD28^+^CD8^+^ T cells. The color represents p.value and the x-axis represents gene count.

### HLA-DR^+^CD38^+^CD8^+^ cluster was characterized by *GZMB* expression

One characteristic of HLA-DR^+^CD38^+^CD8^+^ cluster was the upregulated expression of *GZMB* (Fig. 8A), which plays an important role in T cell cytotoxicity. Metabolic process related signaling pathways were enriched (Fig. 8B), indicating the active status of this cluster.

**Figure 8.**
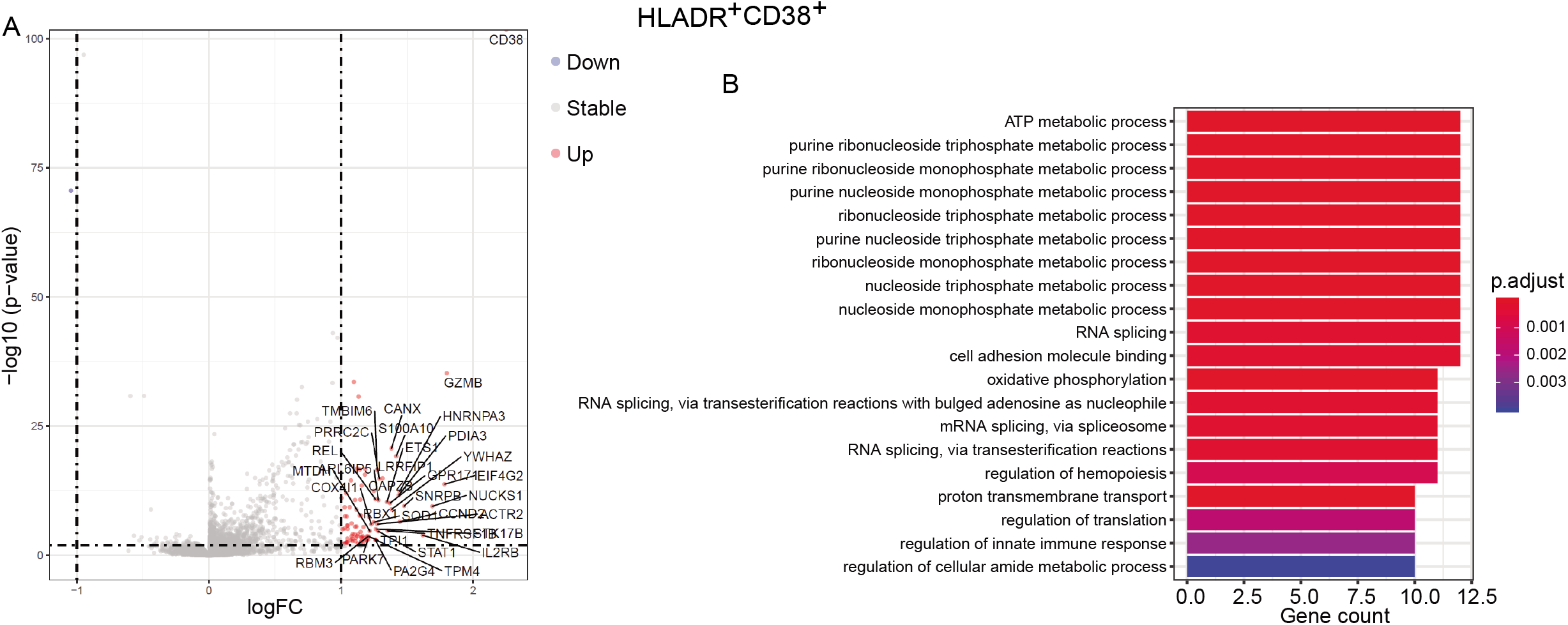
Characteristics of cluster HLA-DR^+^CD38^+^CD8^+^. A, Volcano plot showing differentially expressed genes between HLA-DR^+^CD38^+^ and non-HLA-DR^+^CD38^+^ T cells. Each red or blue dot denotes an individual upregulated or downregulated gene (|logFC| >= 1 and p.value < 0.01). B, Bar plot showing the top 20 pathways of HLA-DR^+^CD38^+^CD8^+^ T cells. The color represents p.value and the x-axis represents gene count.

## Discussion

CX3CR1 expression on CD8^+^ T cells is associated with cytotoxic capability [5, 6]. Consistently, DEG analysis of CX3CR^+^CD8^+^ cluster was characterized by upregulated expression of cytotoxic genes *GZMA* and *NKG7* (Fig. 5A) and T cell activation signaling was top enriched in this cluster (Fig. 5B). But the percentage of CX3CR^+^CD8^+^ cluster was quite low in T cell populations (Fig. 3, Fig. 4), indicating a weak role of this cluster under the circumstances. HLA-DR^+^CD8^+^ T cells are considered activated cytotoxic T lymphocytes [7] and HLA-DR^+^CD28^+^CD8^+^ T cells showed telomerase activity with proliferative potential [8]. The percentages of clusters CX3CR^+^CD8^+^, HLA-DR^+^CD8^+^, and HLA-DR^+^CD28^+^CD8^+^ were downregulated, while the percentage of HLA-DR^+^CD38^+^CD8^+^ cluster which was defined as activated T cells during the acute phase of viral infections [9], was upregulated after antigen stimulation compared to that in unstimulated Ctrl group (Fig. 3). The results indicated various changes of proportions of different cell subsets, though they might have similar functions. On another aspect, the percentages of clusters HLA-DR^+^CD28^+^CD8^+^ and HLA-DR^+^CD38^+^CD8^+^ were decreased with the increased proportion of PD-L1^+^ tumor cells (Fig. 3), implying the inhibition of tumor PD-L1 on the percentages of clusters HLA-DR^+^CD28^+^CD8^+^ and HLA-DR^+^CD38^+^CD8^+^ might result in the inhibition on TCR-T cell cytotoxicity (unpublished data).

The percentages of clusters HLA-DR^+^CD28^+^CD8^+^ and HLA-DR^+^CD38^+^CD8^+^ were higher in TCR-T_MART-1_ than in T_null_, while there was no significant change in the percentages of clusters CX3CR^+^CD8^+^ and HLA-DR^+^CD8^+^ between T_null_ and T_MART-1_ (Fig. 4). It might be the reason why TCR-T_MART-1_ were more cytotoxic than T_null_ (Fig. 2).

## Conclusions

In conclusion, landscape of different functional cell clusters in T_null_ and TCR-T_MART-1_ responding to different proportions of PD-L1^+^ MEL-526 cells loaded with MART-1_27-35_ peptide was provided in this study.

## Materials and methods

### Cell culture

HEK293T (ATCC, CRL-11268) cell line was purchased from ATCC, and MEL-526 (BNCC340404) cell line was purchased from BNCC, and they were cultured in DMEM (Gibco, 21063029) added with 10% fetal bovine serum (Hyclone, SH30084.03HI), penicillin (100 IU/mL), and streptomycin (50 μg/mL) at 37□ and 5% CO_2_. CD8^+^ T cells were cultured in HIPP-T009 (Bioengine, RG0101302) containing 2% fetal bovine serum (Hyclone, SH30084.03HI), IL-2 (20 ng/ml), IL-7 (10 ng/ml) and IL-15 (10 ng/ml) at 37□ and 5% CO_2_.

### Peptide

HLA-A*0201-restricted MART-1 peptide ELAGIGILTV) was synthesized by GenScript (Nanjing, China). Peptide was stored at 10 mg/ml in 100% dimethyl sulfoxide (DMSO; Sigma-Aldrich) at −20°C.

### Plasmid construction

The constant regions of TCR_MART-1_ sequence, which was identified from our previous work (data unpublished), were replaced by mouse TCR constant region α and β, respectively. TCR_MART-1_-encoded DNA was then synthesized by GeneScript (Nanjing, China) and ligated into a lentiviral vector, pRRLSIN.cPPT.PGK (Addgene, 12252).

### Lentivirus production

To perform lentivirus production, 293T cells were transfected with lentiviral vector containing the gene of interest and the packaging constructs (PsPAX2 and PMD2G). The culture medium was collected 72 h after transfection and filtered with 0.45 uM filters (Sartorius). Subsequently, the virus was concentrated by ultracentrifugation at 35,000 rpm for 90 min.

### Generation of MART-1 peptide specific TCR-T cells

Human Peripheral Blood Mononuclear Cells (PBMCs) were isolated from the blood of HLA-A*0201-restricted healthy donors with informed consent. CD8^+^ T cells were purified from PBMC via human CD8 MicroBeads (Miltenyi Biotec) and activated with T Cell TransAct (Miltenyi Biotec); 36-48 h after activation, CD8^+^ T cells were transduced with lentivirus in a 6-well or 12-well plate. To promote infection efficiency, polybrene was added into the medium at the final concentration of 2μg/mL and the well plate was centrifuge at 800g for 30 minutes. T cells were then expanded and maintained in T cell medium.

### In vitro killing assays

TCR-T cells were co-cultured with target cells labeled with Carboxyfluorescein succinimidyl ester (CFSE; Invitrogen) at different E:T ratios for 24 h. Cells were then collected and stained with PI for FACS analysis. The cytotoxicity was calculated with the proportion of PI^+^ CFSE^+^ cells divided by the proportion of CFSE^+^ cells.

### Statistical analysis

PRISM 6 (GraphPad Software) and RStudio were used for data analysis. *P<0.05, **P<0.005, ***P < 0.001. Error bars represented the Mean±SD.

### Differential gene expression analysis

Seurat FindMarkers were used for DEGs analysis. DEGs of each subset were generated relative to all the remained cells. Then DEGs were identified as the criteria: FDR adjusted p value of F test < 0.01.

### Gene set enrichment analysis

The “enrichGO” function in the “clusterProfiler” package was used to perform GO analysis with the corresponding default parameters. Pathways with the q value <0.05 corrected by FDR were used for further analysis.

### Data availability

The data that support the findings of this study have been deposited into CNGB Sequence Archive (CNSA: https://db.cngb.org/cnsa/) of CNGBdb with accession number CNP0001109.

### Ethics approval and consent to participate

The study was approved by the Institutional Review Board on Bioethics and Biosafety of BGI. A written information consent was regularly obtained from all donors.

## Acknowledgements

We sincerely thank the support provided by China National GeneBank. This research was supported by National Natural Science Foundation of China (No. 81903159), Guangdong Provincial Key Laboratory of Genome Read and Write (No. 2017B030301011), the Shenzhen Municipal Government of China Peacock Plan (No. KQTD2015033017150531), and Shenzhen Municipal Government of China (No. 20170731162715261).

## Author contributions

Q.G. designed and supervised the project, wrote and revised the manuscript. S.L. performed the bioinformatic analysis. R.D., H.C. and Q.G. performed the experiments. B.K., Y.G. and X.D. helped with the manuscript revision.

